# Accurate prediction of cell composition, age, smoking consumption and infection serostatus based on blood DNA methylation profiles

**DOI:** 10.1101/456996

**Authors:** Jacob Bergstedt, Alejandra Urrutia, Darragh Duffy, Matthew L. Albert, Lluís Quintana-Murci, Etienne Patin

**Affiliations:** Department of Automatic Control, Lund University, Lund SE-221, Sweden; Center for Translational Science, Institut Pasteur, Paris 75015, France; Laboratory of Dendritic Cell Immunobiology, Department of Immunology, Institut Pasteur, Paris 75015, France; INSERM U1223, France.; Department of Cancer Immunology, Genentech, South San Francisco, California 94080, USA; Unit of Human Evolutionary Genetics, Department of Genomes & Genetics, Institut Pasteur, 75015 Paris, France; CNRS UMR-2000, 75015 Paris, France; Center of Bioinformatics, Biostatistics and Integrative Biology, Institut Pasteur, 75015 Paris, France

**Keywords:** Methylation, Prediction, Elastic net, Cellular composition, EWAS, Age

## Abstract

DNA methylation is a stable epigenetic alteration that plays a key role in cellular differentiation and gene regulation, and that has been proposed to mediate environmental effects on disease risk. Epigenome-wide association studies have identified and replicated associations between methylation sites and several disease conditions, which could serve as biomarkers in predictive medicine and forensics. Nevertheless, heterogeneity in cellular proportions between the compared groups could complicate interpretation. Reference-based cell-type deconvolution methods have proven useful in correcting epigenomic studies for cellular heterogeneity, but they rely on reference libraries of sorted cells and only predict a limited number of cell populations. Here we leverage >850,000 methylation sites included in the MethylationEPIC array and use elastic net regularized and stability selected regression models to predict the circulating levels of 70 blood cell subsets, measured by standardized flow cytometry in 962 healthy donors of western European descent. We show that our predictions, based on a hundred of methylation sites or lower, are less error-prone than other existing methods, and extend the number of cell types that can be accurately predicted. Application of the same methods to age, smoking consumption and several serological responses to pathogen antigens also provide accurate estimations. Together, our study substantially improves predictions of blood cell composition based on methylation profiles, which will be critical in the emerging field of medical epigenomics.

## Introduction

Cellular subtypes that compose organisms derive from various differentiation lineages during development. As stem cells differentiate into more specialized cells, their genome accumulates epigenetic modifications, *i.e.*, stable chemical additions to the DNA that can affect gene expression but do not change the DNA sequence, resulting in cell-specific gene expression. DNA methylation (DNAm), a stable epigenetic mark that refers to the attachment of a methyl group to DNA cytosine, plays a key role in cellular differentiation and gene regulation. Epigenome-wide association studies (EWAS) have searched for DNAm sites that covary with disease conditions or disease-related traits, as these DNAm changes could mediate the effects of environmental perturbations on the transcriptional reprogramming of differentiated cells and, in turn, organismal phenotypes (1, 2). However, interpretation of the results can be problematic, because statistical associations between DNAm and a condition of interest could be due to either a perturbation of the epigenetic properties of a cell subtype that causes the condition, or heterogenetiy in the proportions of differentiated cells caused by the condition (3, 4). For example, because rheumatoid arthritis triggers a change in the granulocyte-to-lymphocyte ratio, an EWAS of this disease identified thousands of associated DNAm sites that became non-significant upon correction for cellular heterogeneity (5). Thus, there is a clear need in the epigenomics field for methods that reliably enumerate cell sub-populations from heterogeneous tissues (6).

Currently, the gold standard approach for cell counting is flow cytometry, a laser-based technology that simultaneously detects several fluorescent-labelled protein markers at a single-cell resolution. However, this approach is labour-intensive and costly, requires skilled practitioners, and its performance is affected by sample degradation. Alternatively, cell composition can be indirectly estimated from gene expression profiles, which are known to be cell-specific (7, 8). These methods, referred to as cellular deconvolution, rely on transcriptional profiles of reference cell populations to predict the cellular composition of sampled cell mixtures, which are also strongly affected by degradation and are difficult to standardize. In a seminal study, Houseman and colleagues used projection methods similar to the ones used for gene expression to estimate blood cell mixture proportions from DNAm profiles (9), a more stable molecular measure. Because DNAm changes are thought to be involved directly in the lineage decision of hematopoietic cells (10, 11), they provide a direct link with blood cell identity. This method, referred to as the ’Houseman method’ or ’Houseman model’, uses DNAm profiles from six sorted cell subtypes as a reference, and assumes that the heterogeneous sample of interest is a mixture of these cells, whose proportions are estimated by projecting the sample matrix on to the reference matrix. The method can estimate the proportion of six major immune cells in blood, using a reference library of 600 CpG sites. Koestler *et al.* proposed a refined reference library, called IDOL (12), achieving better estimation of the six subsets with only 300 CpG sites. Although these methods have been extensively used, they only estimate six major cell subsets, and need at least 300 probes, limiting their usefulness as a tool for adequately controlling confounding in EWAS, and for applications in clinical research.

Here, we build novel parsimonious models for predicting the circulating levels of 70 blood cell subsets measured by flow cytometry in 962 healthy donors of the Milieu Intérieur study (13, 14), based on blood DNAm levels at >850,000 sites (Illumina Methylation EPIC beadchip; (15)). The models are based on two key assumptions: 1) methylation at some sites marks differentiation events that can identify a particular blood cell lineage, and 2) only few methylation probes on the EPIC array mark such differentiation events. The first assumption implies a linear relation between the cell proportion in whole blood and methylation levels at the sites that mark it. We therefore use linear regression models to predict blood cell composition from DNAm levels. The second assumption means that only a small fraction of the probes will actually be predictive. We must therefore look for *sparse* models, which discard many of the included predictors in a data-driven fashion.

We use two approaches to build predictive models of immune cell proportions. The two assumptions mentioned above lead naturally to regularized linear regression models. Therefore, to infer optimal models in terms of prediction accuracy, and to investigate how prediction accuracy depends on the number of predictors, we use the elastic net method (16, 17). Similar models have previously been used for the prediction of age, smoking status, alcohol consumption and educational attainment based on DNAm (18, 19). We believe that elastic net regression will be able to find both the predictors that mark differentiation events for the lineage of the cell, but also the numerous probes that are correlated with such predictors. In addition, we use the more stringent selection technique, *stability selection* (20, 21), to find a minimal stable set of predictors for each proportion. Stability selection selects predictors of each immune cell proportion that are consistently predictive in 100 subsamples of the dataset. We then build predictive models from the stability selected set of predictors using ordinary least squares. Compared with the elastic net, stability selection is more demanding of the predictors it selects. Consequently, it targets probes that mark differentiation events that are the most important for the cell. We therefore explore the biological functions associated with the stability selected probes, to improve knowledge of the epigenetic changes that characterize differentiated immune cells. A similar two-pronged approach is used to predict other conditions and traits collected within the Milieu Intérieur study, including age, smoking, height, BMI, routine chemical and hematological laboratory tests, and the serological responses to antigens of 13 common pathogens (22). Several of these traits have not previously been modelled using all DNAm probes jointly. Our study substantially improves predictions of blood cell composition based on DNA methylation profiles, which will be critical for applications in medical epigenomics, forensics and disease prognosis.

## Results

### Optimization of predictive models

To predict immune cell proportions with optimal accuracy, given our assumption of sparsity and linearity, we use elastic net regularization. It is controlled by two regularization parameters: *λ*, which controls the 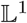 regularization that enforces sparsity on the coefficients, and *α*, which controls 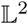 regularization that restricts the magnitude of the coefficients. We use 5 different values for *α* and 200 different values for *λ*. Each possible pair of *α* and *λ* parameter values give a different amount of predictors and regularization, and is a step in the so-called *regularization path*. We measure the prediction accuracy along the regularization path by the mean absolute error (MAE) and the correlation (*R*) between the hold-out sample values and the out-of-sample predictions in 10-fold, twice repeated two-dimensional cross-validation, described in Algorithm 1. The procedure gives 20 samples from the distribution of out-of-sample prediction accuracy along the regularization path. We use those samples to estimate the mean accuracy and its 95% confidence intervals.

The performance of models, together with the number of predictors that is optimal in terms of prediction accuracy, is shown in Table 1 for each cell proportion, as well as 23 other continuous traits, including age and morphometric and physiological measures. DNA methylation levels can accurately predict age and sex (18), intrinsic factors that are predictive of many traits, including immune cell counts in whole blood (14). It is therefore important to discern when predictors based on methylation probes give additional information to these two commonly available factors. For comparison, we therefore include in Table 1 the prediction accuracy of a linear model that only includes age and sex as predictors. We also build predictive models for binary phenotypes, including smoking status and serostatus for 13 different common infections, using elastic net regularization together with the cost function of the binomial likelihood with a logit link function. Similarly to the approach we use for the continuous traits, we estimate prediction accuracy in terms of model complexity using cross-validation. For binary traits we measure prediction accuracy by the classification rate, *i.e.*, the proportion of correct class predictions (probability threshold is taken at 0.5). Prediction accuracy for models with optimally many predictors for the binary traits are shown in Table 2.

**Table 1.**
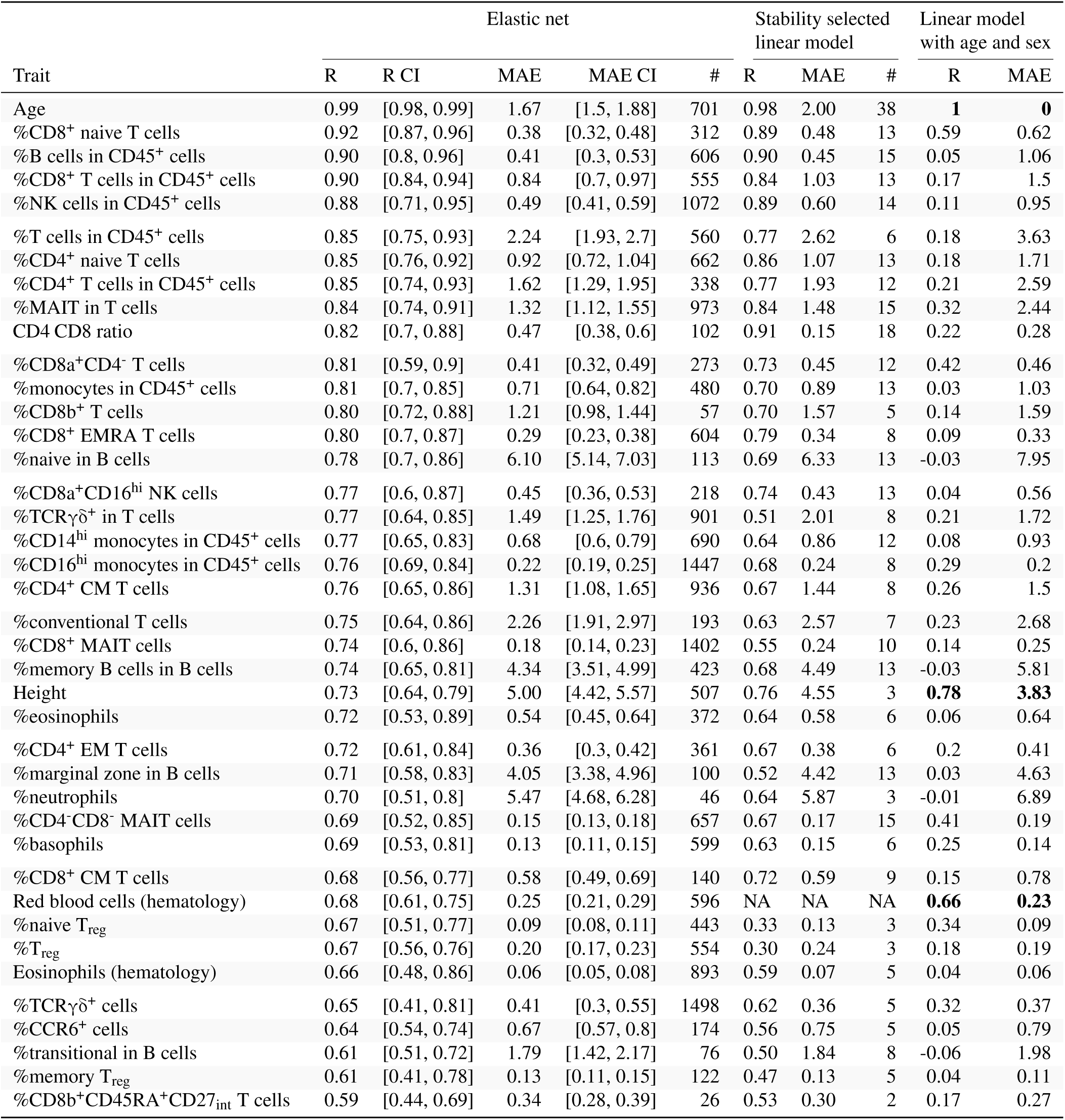
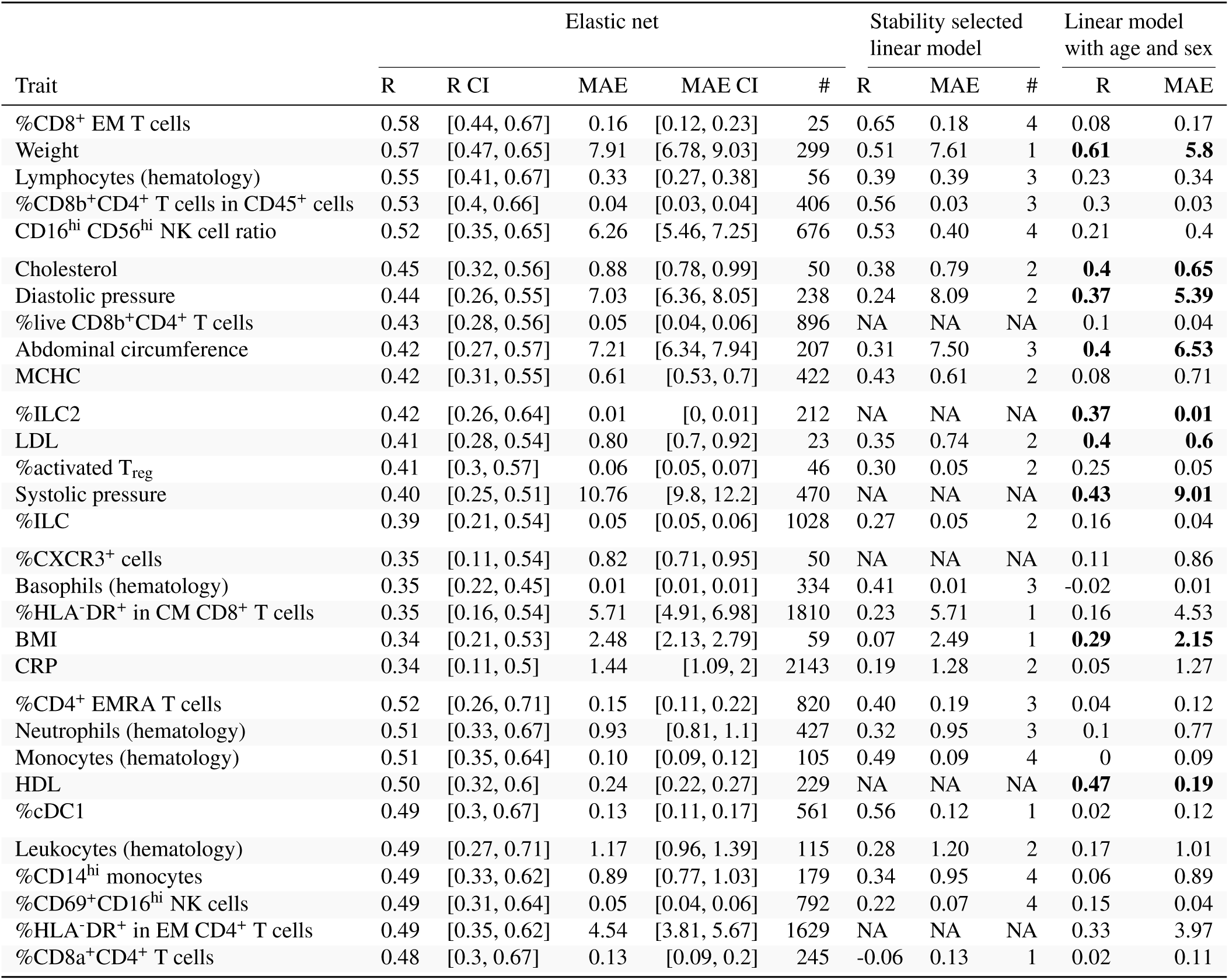
Mean absolute errors (MAE) and correlation (*R*) between out-of-samples predictions and observed values from three different predictive models for each trait. The first results are from elastic net models that have been tuned by our cross-validation scheme, detailed in Algorithm 1. Our scheme gives 20 samples from the distribution of accuracy estimates, which are used to construct confidence intervals. Results are also shown for stability selected linear models. Such models include only predictors that are robustly predictive of the trait. For comparison, the predictive accuracy of each trait is shown also for a simple linear model that only includes age and sex as predictors. Predictive accuracy estimates for the simple model for traits where the difference in *R* between the elastic net model and the simple model is less than 0.10 is shown in bold face. The character ’#’ stands for ’number of probes’. In the case of elastic net models, this is the mean number of probes for the 20 repetitions. *Table continued on next page*.

**Table 2.** Results from the logistic regression elastic net models and stability selected logistic regression models for binary traits. Logistic regression elastic net models were fitted using the logistic regression cost function together with elastic net regularization on the regression parameters. The regularization was tuned using our cross-validation scheme detailed in Algorithm 1. The column ’#’ gives the number of probes included in the model. The column CR gives the classification rate: how many of the out-of-sample classes that were correctly predicted. The naive prediction is to always guess the most prevalent condition. The percentage of people in the whole sample that belongs to the most prevalent class is given in the column “Prev”. Methylation predictors only add something if they can improve on the naive prediction. This measure is given in the “Diff” column which is computed as “CR” - “prevalence”.

| Trait | Prev | Elastic net |  |  |  | Stability selected linear model |  |  |
| --- | --- | --- | --- | --- | --- | --- | --- | --- |
|  |  | CR | CR CI | Diff | # | CR | Diff | # |
| Smoking | 0.52 | 0.89 | [0.82, 0.94] | 0.38 | 192.90 | 0.64 | 0.12 | 3 |
| CMV | 0.65 | 0.87 | [0.81, 0.94] | 0.22 | 255.95 | 0.90 | 0.25 | 13 |
| Toxoplasmosis | 0.56 | 0.72 | [0.65, 0.8] | 0.16 | 1057.65 | 0.69 | 0.13 | 1 |
| Hepatitis B | 0.52 | 0.65 | [0.59, 0.75] | 0.13 | 337.65 | 0.69 | 0.17 | 1 |
| Herpes Simplex 1 | 0.65 | 0.68 | [0.59, 0.76] | 0.03 | 5312.05 | 0.70 | 0.05 | 1 |
| Helicobacter pylori | 0.82 | 0.82 | [0.77, 0.88] | 0.00 | 144.25 | 0.84 | 0.02 | 1 |

### Blood cell deconvolution

Accurate estimations are obtained with elastic net regularized models for 35 immune cell proportions (estimated correlation between predicted and observed out-of-sample values *R*>0.6; Table 1). The four immune cells that we predict with the highest accuracy are CD8^+^ naive T cells, with a correlation between predicted and observed out-of-sample values of *R*=0.92 (95% CI: [0.87, 0.96]), using 312 predictors (95% CI: [295, 338]); B cells (*R*=0.90, 95% CI: [0.8, 0.96]) using 606 predictors (95% CI: [582,635]); CD8^+^ T cells (*R*=0.90,95% CI: [0.84, 0.94]) using 555 predictors (95% CI: [526, 591]); and natural killer (NK) cells (*R*=0.88, 95% CI: [0.71, 0.95]) using 1072 predictors (95% CI: [1036, 1126]). For most immune cell proportions, methylation levels clearly provide additional information in comparison to just age and sex.

A comparison of the performance of our elastic net models and the Houseman model, using either the standard or IDOL reference libraries (9, 12), is given in Table 3. Our models outperform the two models for the six major cell-types that they are currently able to estimate. The correlations between predicted and observed out-of-sample values are systematically higher for our models, relative to the Houseman model with either the default or IDOL reference library (Table 3). Furthermore, our models are less error-prone (Table 3). These findings suggest that elastic net regression models, trained on whole blood standardized cytometry data, can outperform constrained projection techniques based on reference values obtained in a limited number of isolated blood cell sub-types.

**Table 3.** Mean absolute error (MAE) and correlation (*R*) between predicted and observed out-of-sample values compared between our elastic net models and the Houseman model with either the standard reference library or IDOL

| Trait | R |  |  | MAE |  |  |
| --- | --- | --- | --- | --- | --- | --- |
|  | Houseman | IDOL | Elastic net | Houseman | IDOL | Elastic net |
| B cells | 0.875 | 0.853 | 0.898 | 3.043 | 2.752 | 0.408 |
| CD4<sup>+</sup> T cells | 0.783 | 0.824 | 0.847 | 2.207 | 3.522 | 1.619 |
| CD8<sup>+</sup> T cells | 0.824 | 0.852 | 0.896 | 6.368 | 3.768 | 0.840 |
| Monocytes | 0.786 | 0.784 | 0.806 | 3.994 | 2.304 | 0.713 |
| NK cells | 0.788 | 0.813 | 0.884 | 3.512 | 2.232 | 0.494 |
| Neutrophils | 0.634 | 0.634 | 0.703 | 11.377 | 10.751 | 5.471 |

### Linear models selected by stability selection

We next evaluate how prediction accuracy varies with the number of predictors in our models. The regularization paths for the nine best predicted traits are shown in Figure 1. Interestingly, out-of-sample prediction error decreases rapidly with the number of predictors, and plateaus at around 50 predictors (Figure 1). This indicates that accurate predictions can be achieved with much fewer predictors than the hundreds of DNAm probes used by current prediction models of cell composition (9, 12) and age (18, 23). These results suggest that blood cell composition can be predicted well using only a few number of probes that are markers for differentiation events. To find such probes, we estimate a minimal robust predictor set using stability selection. We select and build the models on a subsample of 866 randomly selected individuals, and then evaluate on a hold-out sample of 96 randomly selected individuals.

**Fig. 1.**
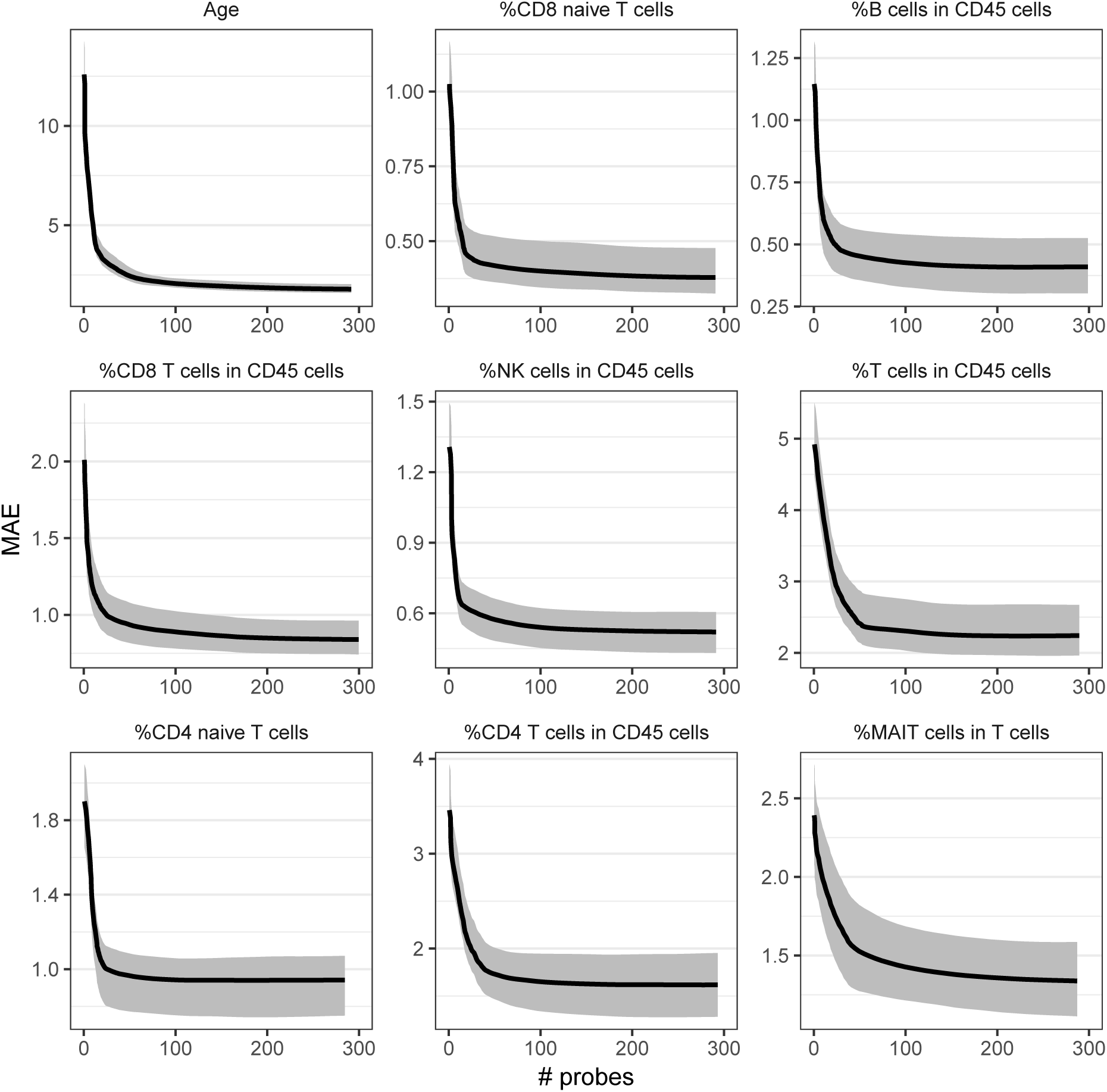
Mean absolute prediction errors (MAE) as a function of the number of predictors included in elastic net models predicting quantitative traits. The regularization parameter *a* is here set to 0.95. Confidence bands are estimated non-parametrically from the 20 samples of the prediction error given by our cross-validation scheme detailed in Algorithm 1.

The predictive accuracy of the stability-selected predictive models is high (Table 1) and comparable to that of elastic net regression models, while using considerably fewer predictors. Prediction performance is also apparent when predicted out-of-sample values are plotted against the observed values for the 16 most accurate models (Figure 2). For instance, using only six methylation probes, the correlation between estimated and observed values for T cells is *R*=0.77 and the MAE is lower than 3%. We verify that our stability selected models are competitive by comparing their prediction accuracy to that of the Houseman model using either the standard or IDOL reference panels. Although our models use only 15, 12, 13, 13, 14 and 3 predictors for B cells, CD4^+^ T cells, CD8^+^ T cells, monocytes, NK cells and neutrophils, respectively, they yield comparable out-of-sample correlations and lower MAE (Table 4), relative to current methods. Together, these results demonstrate that prediction models that use a dozen or fewer methylation probes selected by stability selection can achieve prediction accuracy comparable to that of gold-standard, reference-based cell deconvolution techniques that use hundreds of probes.

**Fig. 2.**
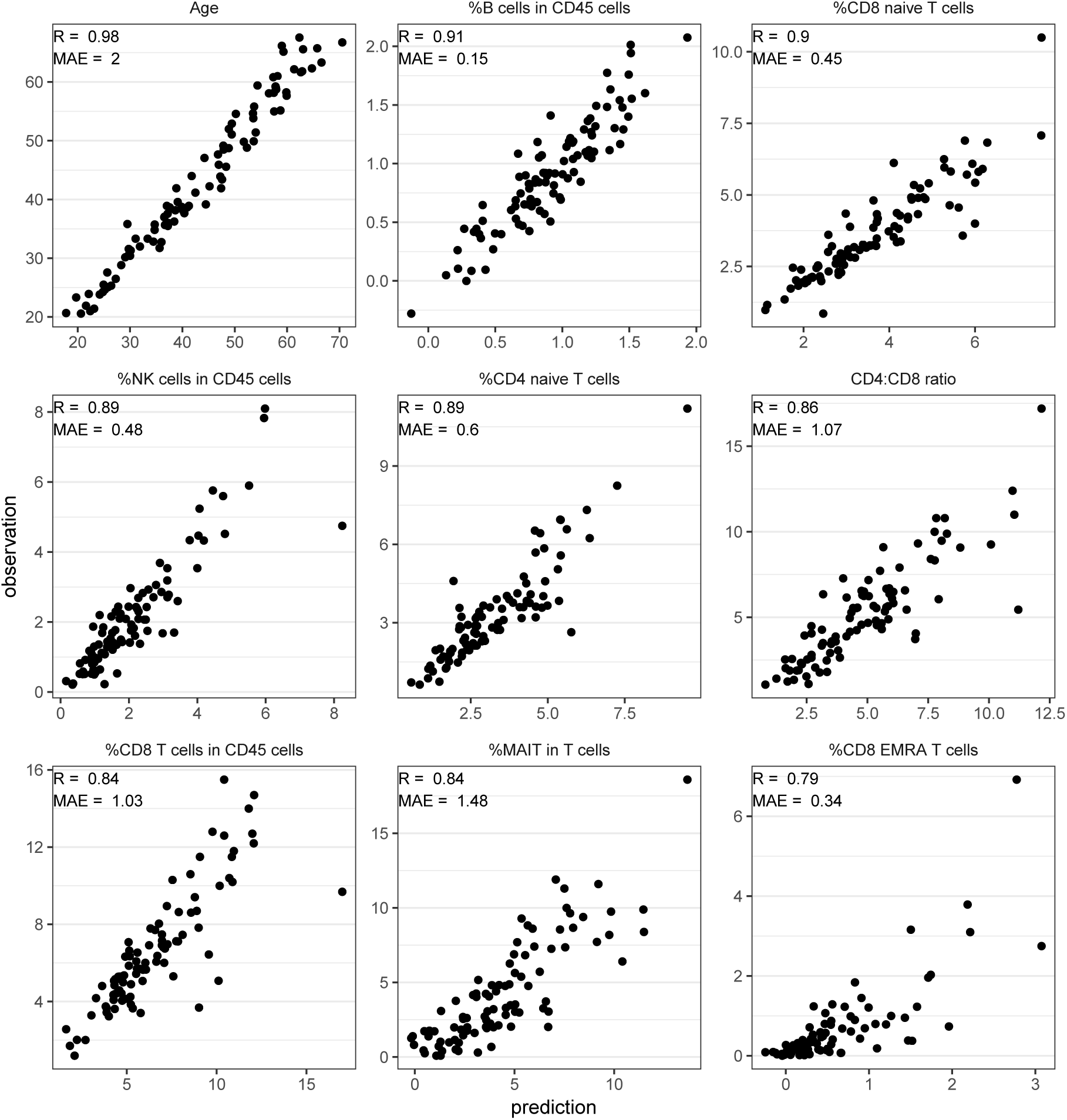
Out-of-sample predictions from stability selected linear models plotted against observed hold-out values. Plots are shown for the 9 best predicted traits. Observed and predicted values are obtained from the hold-out sample of 96 individuals.

**Table 4.** Mean absolute error (MAE) and correlation (*R*) between predicted and observed out-of-sample values compared between our stability selected linear models and the Houseman model with either the standard reference library or IDOL

| Trait | R |  |  | MAE |  |  |
| --- | --- | --- | --- | --- | --- | --- |
|  | Houseman | IDOL | Stab. sel. | Houseman | IDOL | Stab. sel. |
| B cells | 0.899 | 0.889 | 0.905 | 3.107 | 2.846 | 0.454 |
| CD4<sup>+</sup> T cells | 0.757 | 0.788 | 0.774 | 2.622 | 3.645 | 1.933 |
| CD8<sup>+</sup> T cells | 0.829 | 0.825 | 0.844 | 6.381 | 4.024 | 1.032 |
| Monocytes | 0.708 | 0.719 | 0.704 | 3.973 | 2.334 | 0.891 |
| NK cells | 0.799 | 0.802 | 0.891 | 3.591 | 2.259 | 0.600 |
| Neutrophils | 0.609 | 0.609 | 0.640 | 11.333 | 10.666 | 5.871 |

### Biological relevance of the stability selected methylation probes

Because blood cell proportions could be accurately predicted with just a dozen of DNAm probes, we next investigate the relevance of the stability-selected probes to cell biology. We find several, methylome-wide significant DNAm probes that are found close to, or within, genes with well-known functions in immune cell differentiation (Table 5). For instance, DNAm levels within *CD4*, *CD8A* and *CD8B* genes are associated with the CD4:CD8 ratio (*P*=3.9×10^-11^), the proportion of CD8a^+^ NK cells (*P*=4.6×10^-17^) and the proportion of CD8b^+^ T cells (*P*=1.6×10^-9^), respectively. The proportion of neutrophils are associated with DNAm levels in the *PDE4B* gene body (*P*=1.8×10^-8^), which plays a key role in neutrophil function (24). Similarly, the proportion of MAIT cells are associated with DNAm levels in the 5’UTR of *IL21R* (*P*=8.3×10^-21^), which is known to regulate MAIT cell numbers (25). Several cell sub-types, including leukocytes, lymphocytes, monocytes and ILC, are associated with DNAm sites within *AHRR*, *F2RL3* and *GATA3* genes, which are known to be strongly affected by cigarette consumption (26–28). We consistently showed recently that circulating levels of these different blood cell subsets are significantly impacted by smoking status (14). Finally, a number of the selected DNAm probes have previously been associated with disease (Table 5). For instance, DNAm within the *ACSF3* gene is associated with the proportion of naive B cells (*P*=4.1×10^-9^) and has been shown to be differentially methylated in B cells of patients with rheumatoid arthritis (29), suggesting that B cell subtype fractions are altered in these patients. Together, these findings support stability selection as a robust tool to select relevant associated variables, and illustrate the biological relevance of DNAm probes selected as predictors of immune cell proportions.

**Table 5.**
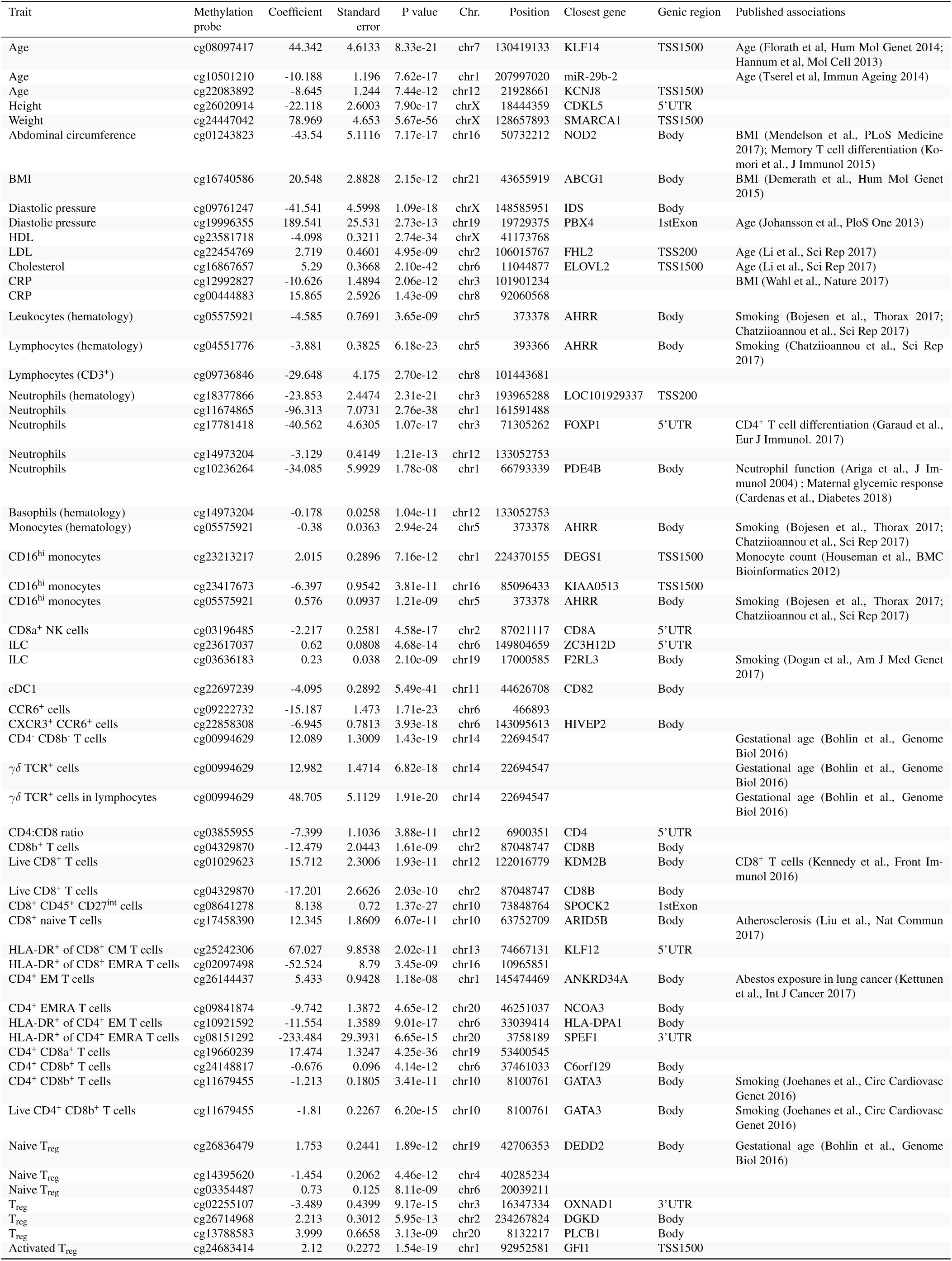
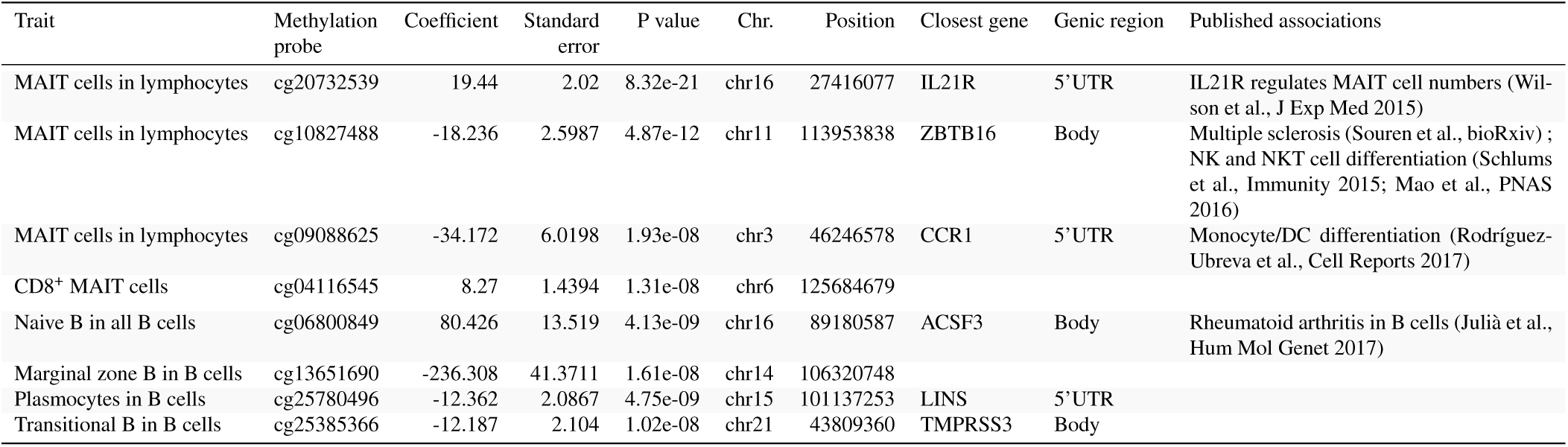
Methylome-wide significant (*P <* 3 × 10^−8^) DNAm probes selected by stability selection for all continuous traits. A model was fitted for each trait with all predictors selected by stability selection. *P* values were then computed for each predictor. The predictors who had *P* values smaller than 3 × 10^−8^ are included in the table. *Table continued on next page*.

### Prediction of other factors

Among the other quantitative factors assessed in the Milieu Intérieur cohort, prediction by the elastic net method is the most accurate for age (Table 1). Using 701 predictors (95% CI: [673, 749]), we estimate age with an MAE of 1.67 years (95% CI: [1.5, 1.88]), confirming that it can be estimated from DNAm with high accuracy (18, 23). From Table 1 it appears that our elastic net models are also able to estimate red blood cell counts, height and weight with high accuracy. However, a comparison with the model that only uses age and sex reveals that the predictive power of methylation levels for these two traits probably mostly stem from their ability to predict age and sex.

We next evaluate the accuracy of elastic net models to predict, based on DNAm data, smoking status and the serostatus for 13 common infections, including infections by *Toxoplasma gondii*, *Helicobacter pylori*, cytomegalovirus (CMV), Epstein-Barr virus (EBV), hepatitis B virus (HBV), Herpes Simplex virus (HSV), Varicella Zoster virus (VZV), mumps virus and measles virus (22). Binary traits for which the prediction of the most prevalent condition outperforms the naive prediction are shown in Table 2. We obtain good prediction results for smoking consumption and CMV serostatus, which is natural considering that both factors have been shown to broadly affect immune cell variation (14). The out-of-sample classifications for both of these traits are correct almost 90% of the time. The estimated regularization paths for the different binary traits are shown in Figure 3. which indicate that near optimal prediction can be achieved with less than 50 DNAm probes. The optimal classification rate for CMV serostatus is *CR*=87% (95% CI: [81%, 94%]) using 256 predictors, while for smoking, *CR*=89% (CI: [82%, 94%]) using 193 predictors.

**Fig. 3.**
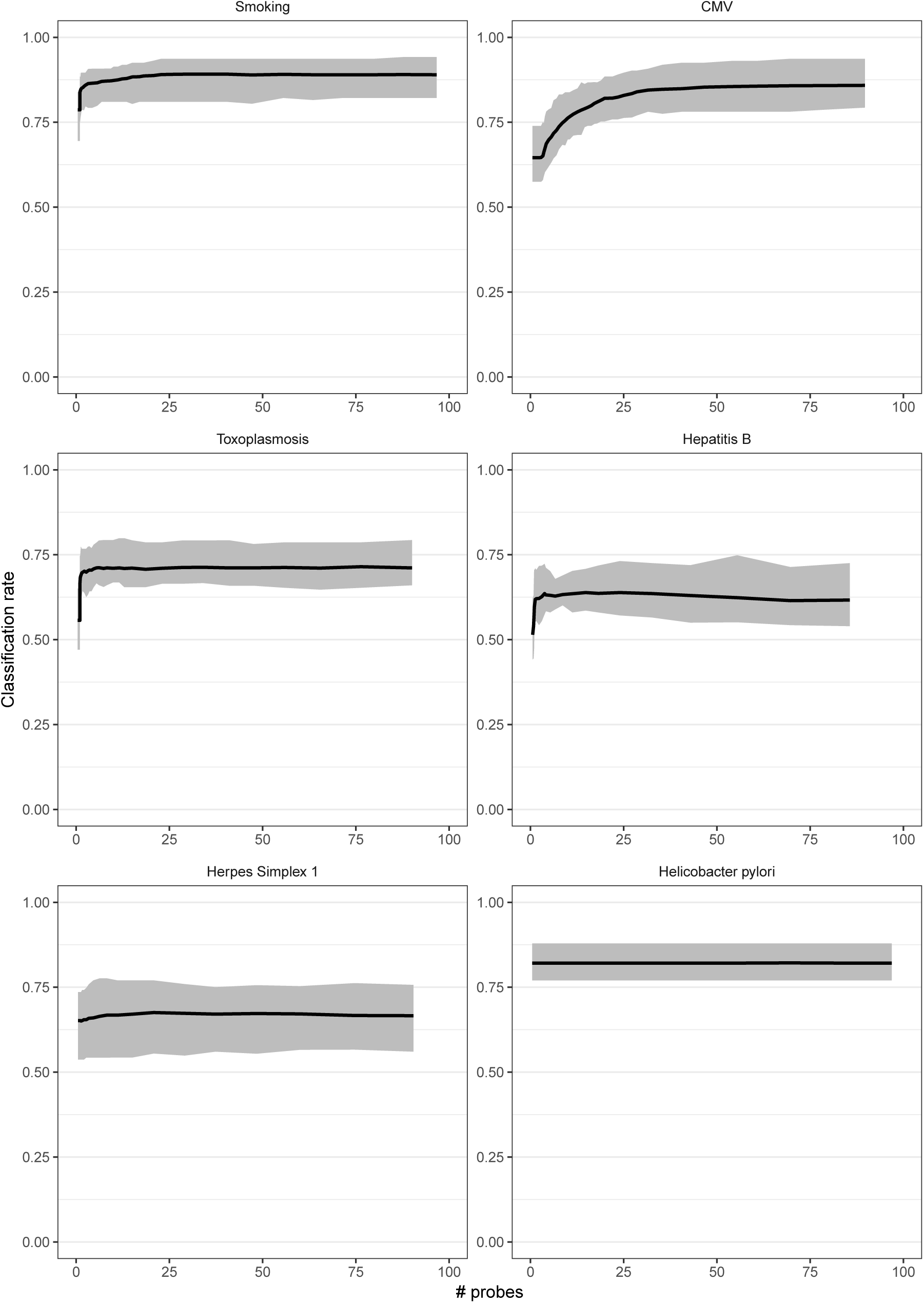
Classifcation rate as a function of the number of predictors included in logistic regression elastic net models predicting binary traits. The regularization parameter *a* is here set to 0.95. The 6 binary traits best predicted by the models are shown. Confdence bands are estimated non-parametrically using the 20 samples from the distribution of predictive accuracy given by our cross-validation scheme detailed in Algorithm 1

We also select a robust minimal set of predictors using stability selection for the binary phenotypes. Models are selected and fitted using the same training set of 866 samples as for continuous traits, and then evaluated on the 96 holdout samples. The prediction accuracy of the models is shown in Table 2. Interestingly, the stability selected model for CMV performs slightly better than the elastic net model, using only 13 probes, while the selected model for smoking performs notably worse. This indicates that the relationship between DNAm and smoking is less sparse than that for DNAm and CMV serostatus. Methylome-wide significant probes selected for smoking are well known DNAm sites predictive of cigarette consumption (Table 6). We find that HBV, *T. gondii* and HVS1 infections associate with DNAm sites close to *EVOLV2* and *KLF14* genes, known to be strongly associated with age. This suggests no effects of these infections on DNAm besides that of age, with which they are themselves strongly correlated (22). More interestingly, DNAm associated with *H. pylori* seropositivity is found within the poliovirus receptor-like 3 gene (*P*=4.6×10^-12^), an intestinal epithelium receptor for bacterial toxins (30), suggesting a role of this protein in *H. pylori* infection.

**Table 6.**
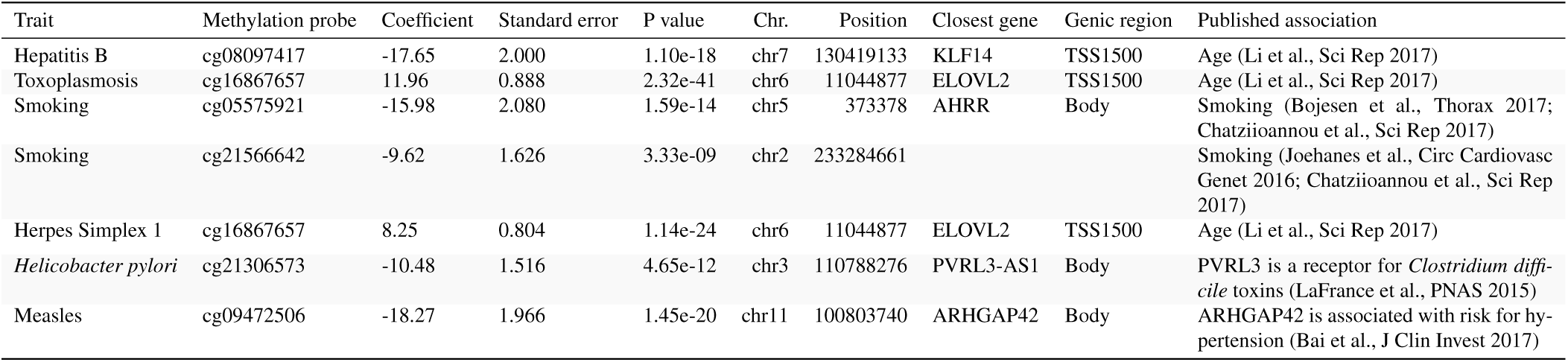
Methylome-wide significant (*P <* 3 × 10^−8^) DNAm probes selected by stability selection for all binary traits. A logistic regression model with all predictors chosen by stability selection was fitted for each trait. The predictors who had p values smaller than 3 × 10^−8^ are included in the table.

| Trait | Methylation probe | Coefficient | Standard error | P value | Chr. | Position | Closest gene | Genic region | Published association |
| --- | --- | --- | --- | --- | --- | --- | --- | --- | --- |
| Hepatitis B | cg08097417 | -17.65 | 2.000 | 1.10e-18 | chr7 | 130419133 | KLF14 | TSS1500 | Age (Li et al., Sci Rep 2017) |
| Toxoplasmosis | cg16867657 | 11.96 | 0.888 | 2.32e-41 | chr6 | 11044877 | ELOVL2 | TSS1500 | Age (Li et al., Sci Rep 2017) |
| Smoking | cg05575921 | -15.98 | 2.080 | 1.59e-14 | chr5 | 373378 | AHRR | Body | Smoking (Bojesen et al., Thorax 2017; Chatzioannou et al., Sci Rep 2017) |
| Smoking | cg21566642 | -9.62 | 1.626 | 3.33e-09 | chr2 | 233284661 |  |  | Smoking (Joehanes et al., Circ Cardiovasc Genet 2016; Chatzioannou et al., Sci Rep 2017) |
| Herpes Simplex 1 | cg16867657 | 8.25 | 0.804 | 1.14e-24 | chr6 | 11044877 | ELOVL2 | TSS1500 | Age (Li et al., Sci Rep 2017) |
| *Helicobacter pylori* | cg21306573 | -10.48 | 1.516 | 4.65e-12 | chr3 | 110788276 | PVRL3-AS1 | Body | PVRL3 is a receptor for *Clostridium difficile* toxins (LaFrance et al., PNAS 2015) |
| Measles | cg09472506 | -18.27 | 1.966 | 1.45e-20 | chr11 | 100803740 | ARHGAP42 | Body | ARHGAP42 is associated with risk for hypertension (Bai et al., J Clin Invest 2017) |

## Discussion

Our study reports novel, accurate models to predict blood cell composition from whole blood DNAm profiles. Models were built using a unique dataset that comprises both the quantification of 70 blood cell proportions by standardized flow cytometry (14) and blood methylomes established with the MethylationEPIC array (15), assessed in 962 healthy donors of western European ancestry. Predictive models are built using the elastic net method (17), a regularized linear regression model that has been recently used to predict age from MethylationEPIC array data (18). The prediction accuracy, measured as the correlation between predicted and observed out-of-sample values and the MAE, is improved for our models, compared to the widely-used Houseman model, based on either the standard or improved IDOL reference libraries (9, 12). We are also able to accurately predict 35 subset frequencies, in contrast to the six that are currently possible to estimate by the Houseman model using either reference panel. These results suggest that our models should better prevent false positives in EWAS due to cellular heterogeneity, relative to existing gold-standard methods. Nevertheless, it must be noted that we assessed prediction accuracy based on cellular fractions estimated with the same flow cytometry technique, panel design and standardization steps as those used for the training dataset, which may disfavor the other methods trained on other types of cell enumeration techniques.

We also show that it is possible to find predictive models of immune cell proportions that are comparable in terms of accuracy to elastic net models, and to the Houseman models with either reference library, using considerably fewer predictors. This is done by employing the stability selection technique (20, 21). Because of their much smaller size, such models can more robustly, flexibly and cost-effectively predict blood cell composition, age, and smoking consumption than previous models.

Thanks to the exhaustive immunophenotyping performed in our training dataset, we can extend the number of blood cell subsets that can be accurately predicted from blood DNAm data. Notably, our models can accurately predict the blood frequencies of MAIT cells, eosinophils, basophils and T_reg_ cells (*R*>0.6; Table 1). Importantly, all these leukocyte subsets have previously been reported to vary with various disease conditions, and are thus expected to confound interpretation of EWAS. For instance, circulating levels of MAIT cells are known to be strongly altered during infection (31) and in systemic lupus erythematosus and rheumatoid arthritis patients (32). Eosinophil numbers change with exposure to allergens and in asthmatic patients (33). Similarly, T_reg_ populations and sub-populations show altered frequencies in several autoimmune and allergic diseases (34). Therefore, adjusting for these newly-predicted cell populations may improve correction for cellular heterogeneity in epigenomic studies of immune-related disorders. More generally, we envisage that prediction models of blood cellular composition could also be employed to better understand disease pathophysiology *per se*. While EWAS assume that disease-associated DNAm sites affect the transcriptional reprogramming of already differentiated cells, there is increasing evidence that diseases can also be caused by stable alterations of cellular repertoires, a phenomenon recently referred to as polycreodism (4). We suggest that model-based estimation of blood cell composition in large longitudinal cohorts, for which methylomes but no flow cytometric measurements exist, will represent a powerful new approach to evaluate whether perturbations in cell proportions can predict disease outcome.

## Methods

### DNA methylation data

The *Milieu Intérieur* cohort includes 1,000 healthy donors who were recruited by BioTrial (Rennes, France) and were stratified by gender (*i.e.*, 500 women and 500 men) and age (*i.e.*, 200 individuals from each decade of life, between 20 and 70 years of age). Donors were selected based on stringent inclusion and exclusion criteria, detailed elsewhere (13). DNAm data was retrieved for all donors from a previous study (15), where detailed methods are provided. In brief, the DNA methylome was profiled with the Infinium MethylationEPIC BeadChip on whole blood-derived samples. Raw fluorescence intensities of 866,895 methylation sites across the human genome were processed with the *R* (version 3.5) *Bioconductor* package *minfi*. Values were corrected for probe color bias and differences in type-I and type-II probe distributions, using the single sample NOOB (ssNOOB) method implemented in minfi. Because we wanted to use the methylation data primarily for prediction, which can easily be evaluated on out-of-sample observations and in validation cohorts, we wanted to exclude as few probes as possible. Therefore, we did not exclude probes from the X and Y chromosomes. We did neither exclude possibly cross-reactive probes. From the 866,895 initial probes, we only excluded probes that had a *detection P* ≥ 0.01 for *more* than 3 samples. A total of 858,923 probes were kept for the analyses. We suppose in this study that DNAm levels are linearly related to cell proportions. We therefore use β methylation values instead of *m* values.

### Flow cytometry data

Flow cytometry data was retrieved for all *Milieu Intérieur* donors from a previous study (14), where detailed methods are provided. Briefly, whole blood samples were collected from the 1,000 healthy, fasting donors on Li-heparin. Sample staining was performed within 6h of blood draw.Ten 8-color flow cytometry panels were developed. The acquisition of cells was performed using two MACSQuant analyzers, which were calibrated using MacsQuant calibration beads. Flow cytometry data were generated using MACSQuantify™ software. Among the 313 exported immunophenotypes, we only kept 70 cell proportions and 2 ratios as candidate measures for prediction.

### Houseman model using standard and IDOL reference libraries

We used the implementation of the Houseman model in the *EstimateCellCounts2* function of the *Bioconductor R* package *FlowSorted.Blood.EPIC* to predict immune cell proportions for all our 962 samples with both the default and IDOL reference panels.

### Statistical modeling

We suppose that there are DNAm CpG sites in the genome of a cell that mark a particular cellular lineage, in the sense that the methylation state of these sites are specific to the cells belonging to that lineage. Therefore, we expect the state of methylation at a number of CpG sites to mark the identity of a particular blood cell. In whole blood, the percentage of cells that are methylated at such DNAm sites should be linearly related to the proportion of the cell in the blood. We further suppose that it is primarily such DNAm sites, and sites related to them, that are predictive of blood cell proportions. We therefore use a linear model to predict blood cell proportions from DNA methylation levels in whole blood. Let 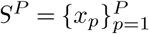*, P* = 858923 denote our observations of the percentage of methylation at all measured DNAm sites. Let *C* ≪ *P* be the number of sites that are related to a differentiation event that offers information on the identity of a particular blood cell. This could be a primary event that directly determines cell identity, or it could be an event that gives information on the identity of the cell because of the correlation structure with other cells or genetic and environmental factors. We expect that only few sites correspond to primary events and we further expect the average methylation at such sites to be highly predictive of the immune cell proportion whose lineage it marks. We expect more events that offer correlational information on the proportion of immune cells. Typically, such sites are distributed according to a long tail of decreasing predictive power. Let *D* ≪ *C* be the number of sites that correspond to primary differentiation events for a particular blood cell. To summarize: for a particular blood cell, we are targeting two sets of probes, *S^C^* and *S^D^* such that

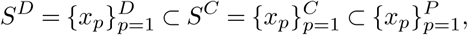

and we suppose a predominantly linear relationship between these variables and the cell proportion. We are therefore looking for sparse linear models, where the coefficients of the predictors in *S^P^* \ *S^C^* is set to zero. We employ two different strategies to target the predictors in *SC* and *SD*. Let *n* = 962 be our sample size. We expect *D* ≪ *n*, but do not necessarily suppose that *C < n*. For *S^C^* we therefore need to *select* predictors, but the linear regression equation system could still be overdetermined, so we also need to *regularize* the coefficients of the fitted linear model. To do this, we employ elastic net regularization (17). In the case of *S^D^*, we only want to *select* predictors and then fit an unbiased least squares regression model. We achieve this by using the stability selection technique (20, 21).

### Elastic net regression

Let now 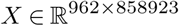 be the matrix corresponding to *S^P^*, with the methylation percentages as columns. Furthermore, let 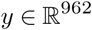 be observations of a cell proportion. The elastic net cost function combines the least squares term with two regularization terms on the magnitude of the coefficients for the columns in *X*
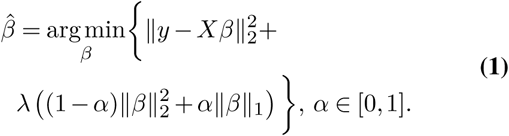

The parameter *α* chooses between a pure Euclidian norm squared penalty, 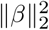, corresponding to ridge regression at *α* = 0 and a pure 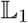 norm, ∥*β*∥_1_, penalty corresponding to the LASSO (16) penalty at *α* = 1. If *α* ≠ 0 then the estimator in Eq. (1) will do a selection: coefficients that do not rise above a noise floor will be put to exactly zero. The pure LASSO penalty has a *saturation* property: it cannot select more predictors than the number of samples (35). Note that for the pure LASSO penalty, all coefficients will be zero if
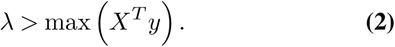

To target *X^C^*, we suppose that an *α* between zero and one will be optimal. To find this parameter, we employ our own cross-validation scheme, detailed in Algorithm 1. We fit the optimization problem Eq. (1) by the *glmnet* package in *R*.

### Stability selected linear regression

Elastic net regression with regularization parameters tuned by cross-validation will typically include predictors of weak predictive power as well as some false positives (36). To target *SD*, we therefore use a more stringent selection scheme. As mentioned above, we suppose that *D* ≪ *n*. Therefore, we are now only aiming to select predictors to use in a linear model; we do not want to regularize the parameters. Define the *support S* of a linear model by

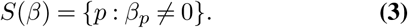

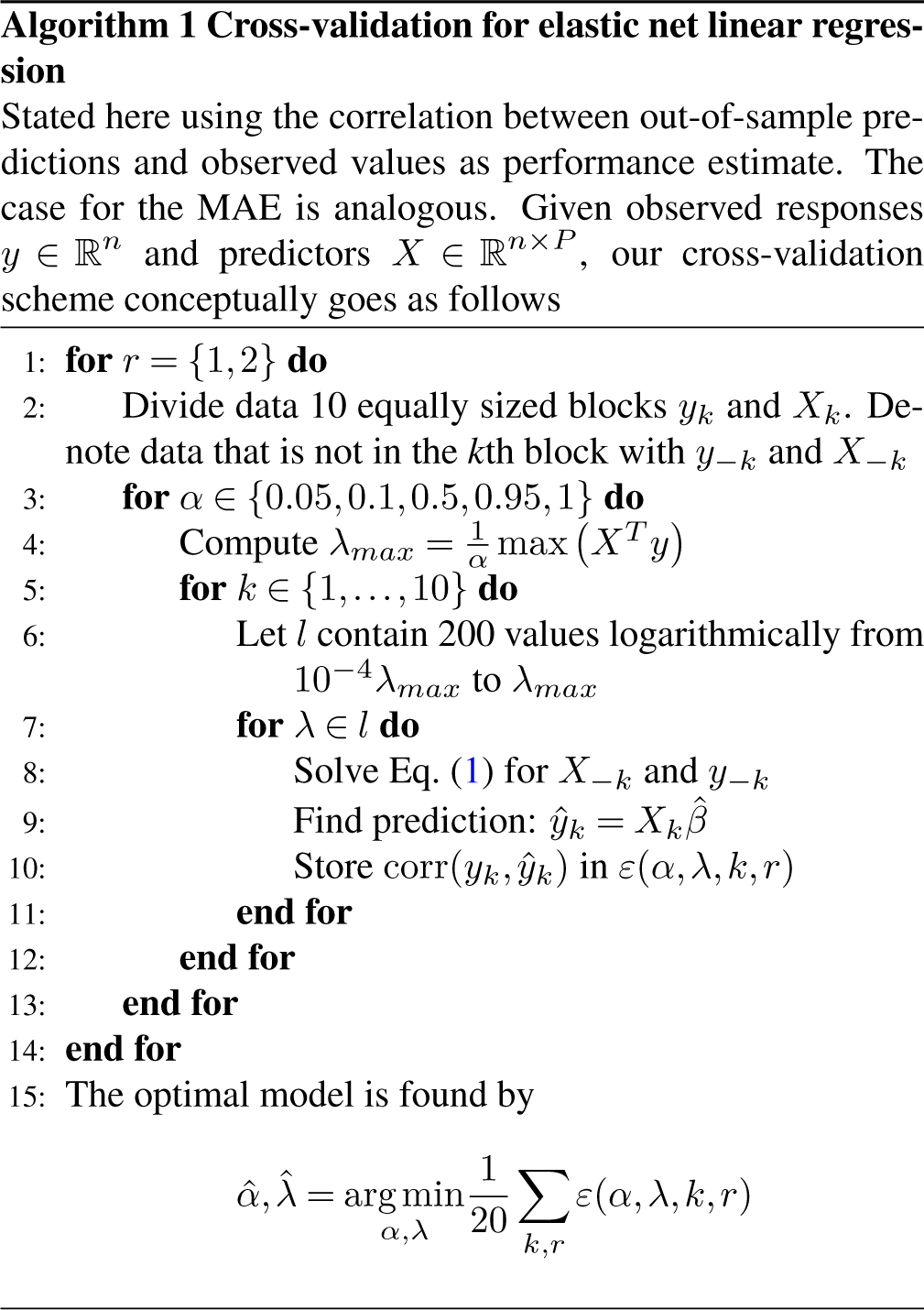

First we introduce a weak support estimator. This estimator uses the cost function in Eq. (1) with *α* fixed at 0.8, while keeping *λ* large enough so that it never includes more than *q* variables. Given this constraint, the support is then estimated to be the included variables. To be more precise, introduce the family of support estimators

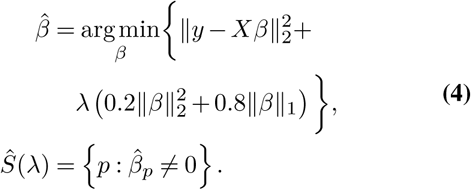

We then use the support estimator *Ŝ_q_* = *Ŝ*(*λ*^*^), where *λ*^*^ is such that

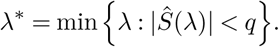

To find *S^D^*, we wrap this weak support estimator in a sub-sampling scheme known as stability selection. The full scheme is outlined in Algorithm 2. Let *X_ss_* be the columns of *X* corresponding to predictors selected by the stability selection scheme. The coefficient estimates for the final linear regression model of the immune cell proportion with measurements in *y* is then
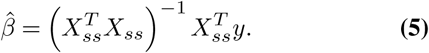

We use the implementation of stability selection in the *stabs R* package (37).

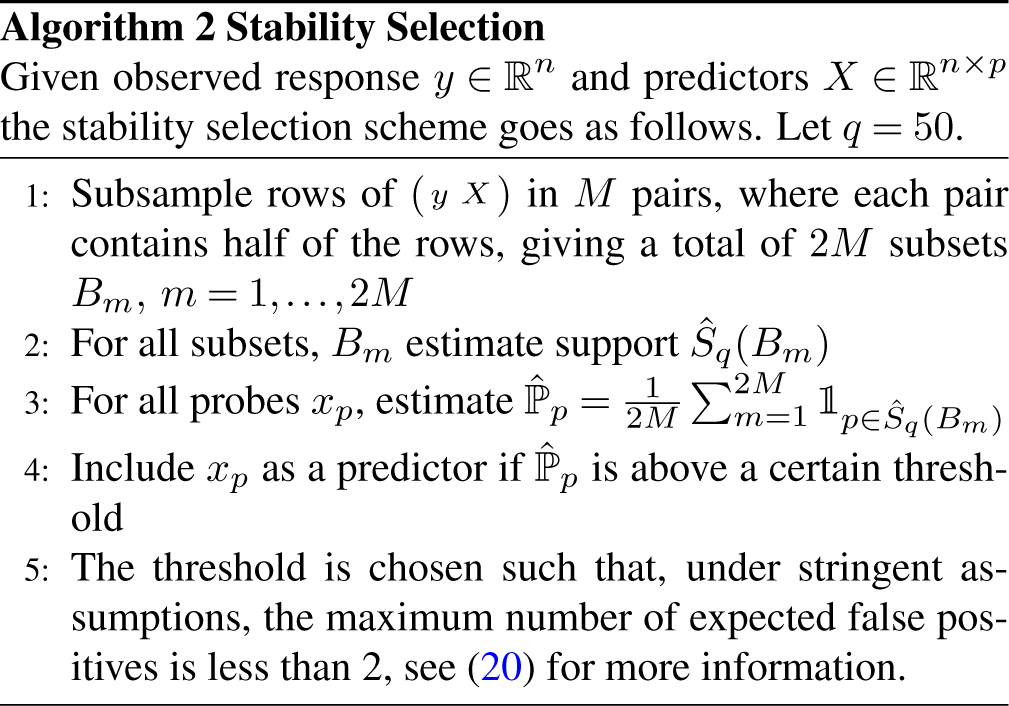

### Other traits

The models above were developed primarily for immune cell proportions, but we use them also for the other traits. We suppose that most of the predictive power of whole blood DNA methylation for any trait comes from its intimate link with immune cell proportions. Therefore, we anticipate that prediction models of a form suitable for immune cell frequencies should work well also for traits related to them.

For binary traits, code the classes as either 0 or 1. The procedure we use for binary traits follows the algorithms above verbatim, except that the least squares term 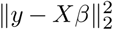 in the cost function in Eq. (1) is replaced by the negative log-likelihood of the binomial distribution given a logit link function

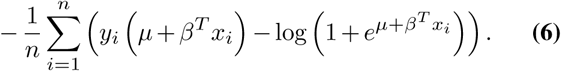

Logistic regression with elastic net regularization is implemented in *glmnet*. For stability selection, we use the *stabs R* package with a custom built selection function based on *glmnet*.

## ACKNOWLEDGEMENTS

This work benefited from support of the French government’s Program *Investissement d’Avenir*, managed by the Agence Nationale de la Recherche (ANR, reference 10-LABX-69-01). J.B. is a member of the LCCC Linnaeus Center and the ELLIIT Excellence Center at Lund University and is supported by the ELLIIT Excellence Center.

## Bibliography

1. Randy L Jirtle and Michael K Skinner. Environmental epigenomics and disease susceptibility. Nature Reviews Genetics, 8(4):253, 2007.

2. Robert Feil and Mario F Fraga. Epigenetics and the environment: emerging patterns and implications. Nature Reviews Genetics, 13(2):97, 2012.

3. Vardhman K Rakyan, Thomas A Down, David J Balding, and Stephan Beck. Epigenomewide association studies for common human diseases. Nature Reviews Genetics, 12(8): 529, 2011.

4. Tuuli Lappalainen and John M Greally. Associating cellular epigenetic models with human phenotypes. Nature Reviews Genetics, 18(7):441, 2017.

5. Yun Liu, Martin J Aryee, Leonid Padyukov, M Daniele Fallin, Espen Hesselberg, Arni Runarsson, Lovisa Reinius, Nathalie Acevedo, Margaret Taub, Marcus Ronninger, et al. Epigenome-wide association data implicate dna methylation as an intermediary of genetic risk in rheumatoid arthritis. Nature biotechnology, 31(2):142, 2013.

6. Andrew E Teschendorff and Caroline L Relton. Statistical and integrative system-level analysis of dna methylation data. Nature Reviews Genetics, 19(3):129, 2018.

7. Aaron M Newman, Chih Long Liu, Michael R Green, Andrew J Gentles, Weiguo Feng, Yue Xu, Chuong D Hoang, Maximilian Diehn, and Ash A Alizadeh. Robust enumeration of cell subsets from tissue expression profiles. Nature Methods, 12(5):453, 2015.

8. Shai S Shen-Orr and Renaud Gaujoux. Computational deconvolution: extracting cell type-specifc information from heterogeneous samples. Current Opinion in Immunology, 25(5): 571–578, 2013.

9. Eugene Andres Houseman, William P Accomando, Devin C Koestler, Brock C Christensen, Carmen J Marsit, Heather H Nelson, John K Wiencke, and Karl T Kelsey. Dna methylation arrays as surrogate measures of cell mixture distribution. BMC Bioinformatics, 13(1):86, 2012.

10. Damiana Álvarez-Errico, Roser Vento-Tormo, Michael Sieweke, and Esteban Ballestar. Epigenetic control of myeloid cell differentiation, identity and function. Nature Reviews Immunology, 15(1):7, 2015.

11. Aimée M Deaton, Shaun Webb, Alastair RW Kerr, Robert S Illingworth, Jacky Guy, Robert Andrews, and Adrian Bird. Cell type–specific dna methylation at intragenic cpg islands in the immune system. Genome Research, 2011.

12. Devin C Koestler, Meaghan J Jones, Joseph Usset, Brock C Christensen, Rondi A Butler, Michael S Kobor, John K Wiencke, and Karl T Kelsey. Improving cell mixture deconvolution by id entifying o ptimal dna methylation l ibraries (idol). BMC Bioinformatics, 17(1):120, 2016.

13. Stéphanie Thomas, Vincent Rouilly, Etienne Patin, Cécile Alanio, Annick Dubois, Cécile Delval, Louis-Guillaume Marquier, Nicolas Fauchoux, Seloua Sayegrih, Muriel Vray, Darragh Duffy, Lluis Quintana-Murci, and Matthew L. Albert. The milieu intérieur study — an integrative approach for study of human immunological variance. Clinical Immunology, 157 (2):277–293, 2015. ISSN 1521-6616. doi: https://doi.org/10.1016/j.clim.2014.12.004.

14. Etienne Patin, Milena Hasan, Jacob Bergstedt, Vincent Rouilly, Valentina Libri, Alejandra Urrutia, Cécile Alanio, Petar Scepanovic, Christian Hammer, Friederike Jönsson, Benoît Beitz, Hélène Quach, Yoong Wearn Lim, Julie Hunkapiller, Magge Zepeda, Cherie Green, Barbara Piasecka, Claire Leloup, Lars Rogge, François Huetz, Isabelle Peguillet, Olivier Lantz, Magnus Fontes, James P. Santo, Stéphanie Thomas, Jacques Fellay, Darragh Duffy, Lluís Quintana-Murci, and Matthew L. Albert. Natural variation in the parameters of innate immune cells is preferentially driven by genetic factors. Nature Immunology, 19(3):302–314, 2018. ISSN 1529-2916. doi: 10.1038/s41590-018-0049-7.

15. Sadoune Ait Kaci Azzou, Etienne Patin, Alejandra Urrutia, Hélène Quach, Jacob Bergstedt, Darragh Duffy, Michael S Kobor, Matthew L Albert, and Lluís Quintana-Murci. Limited impact of environmental exposures on the human blood methylome in adulthood. in preparation.

16. Robert Tibshirani. Regression shrinkage and selection via the lasso. Journal of the Royal Statistical Society. Series B (Statistical Methodology), pages 267–288, 1996.

17. Hui Zou and Trevor Hastie. Regularization and variable selection via the elastic net. Journal of the Royal Statistical Society: Series B (Statistical Methodology), 67(2):301–320, 2005.

18. Qian Zhang, Costanza Vallerga, Rosie Walker, Tian Lin, Anjali Henders, Grant Montgomery, Ji He, Dongsehng Fan, Javed Fowdar, Martin Kennedy, Toni Pitcher, John Pearson, Glenda Halliday, John Kwok, Ian Hickie, Simon Lewis, Tim Anderson, Peter Silburn, George Mellick, Sarah E Harris, Paul Redmond, Alison Murray, David Porteous, Christopher Haley, Kathryn Evans, Andrew McIntosh, Jian Yang, Jacob Gratten, Riccardo Marioni, Naomi Wray, Ian Deary, Allan Mcrae, and Peter Visscher. Improved prediction of chronological age from dna methylation limits it as a biomarker of ageing. bioRxiv, 2018. doi: 10.1101/327890.

19. Daniel L McCartney, Anna J Stevenson, Stuart J Ritchie, Rosie M Walker, Qian Zhang, Stewart W Morris, Archie Campbell, Alison D Murray, Heather C Whalley, Catharine R Gale, David J Porteous, Chris S Haley, Allan F McRae, Naomi R Wray, Peter M Visscher, Andrew M McIntosh, Kathryn L Evans, Ian J Deary, and Riccardo E Marioni. Epigenetic prediction of complex traits and death. bioRxiv, 2018. doi: 10.1101/294116.

20. Rajen D Shah and Richard J Samworth. Variable selection with error control: another look at stability selection. Journal of the Royal Statistical Society: Series B (Statistical Methodology), 75(1):55–80, 2013.

21. Nicolai Meinshausen and Peter Bühlmann. Stability selection. Journal of the Royal Statistical Society: Series B (Statistical Methodology), 72(4):417–473, 2010.

22. Petar Scepanovic, Cécile Alanio, Christian Hammer, Flavia Hodel, Jacob Bergstedt, Etienne Patin, Christian W Thorball, Nimisha Chaturvedi, Bruno Charbit, Laurent Abel, et al. Human genetic variants and age are the strongest predictors of humoral immune responses to common pathogens and vaccines. Genome Medicine, 10(1):59, 2018.

23. Steve Horvath. Dna methylation age of human tissues and cell types. Genome Biology, 14 (10):3156, 2013.

24. Miyako Ariga, Barbara Neitzert, Susumu Nakae, Genevieve Mottin, Claude Bertrand, Marie Pierre Pruniaux, S-L Catherine Jin, and Marco Conti. Nonredundant function of phosphodiesterases 4d and 4b in neutrophil recruitment to the site of inflammation. The Journal of Immunology, 173(12):7531–7538, 2004.

25. Robert P Wilson, Megan L Ives, Geetha Rao, Anthony Lau, Kathryn Payne, Masao Kobayashi, Peter D Arkwright, Jane Peake, Melanie Wong, Stephen Adelstein, et al. Stat3 is a critical cell-intrinsic regulator of human unconventional t cell numbers and function. Journal of Experimental Medicine, 212(6):855–864, 2015.

26. Roby Joehanes, Allan C Just, Riccardo E Marioni, Luke C Pilling, Lindsay M Reynolds, Pooja R Mandaviya, Weihua Guan, Tao Xu, Cathy E Elks, Stella Aslibekyan, et al. Epigenetic signatures of cigarette smoking. Circulation: Genomic and Precision Medicine, 9(5): 436–447, 2016.

27. Stig E Bojesen, Nicholas Timpson, Caroline Relton, George Davey Smith, and Børge G Nordestgaard. Ahrr (cg05575921) hypomethylation marks smoking behaviour, morbidity and mortality. Thorax, 72(7):646–653, 2017.

28. Aristotelis Chatziioannou, Panagiotis Georgiadis, Dennie G Hebels, Irene Liampa, Ioannis Valavanis, Ingvar A Bergdahl, Anders Johansson, Domenico Palli, Marc Chadeau-Hyam, Alexandros P Siskos, et al. Blood-based omic profiling supports female susceptibility to tobacco smoke-induced cardiovascular diseases. Scientific Reports, 7:42870, 2017.

29. Antonio Julià, Devin Absher, María López-Lasanta, Nuria Palau, Andrea Pluma, Lindsay Waite Jones, John R Glossop, William E Farrell, Richard M Myers, and Sara Marsal. Epigenome-wide association study of rheumatoid arthritis identifies differentially methylated loci in b cells. Human Molecular Genetics, 26(14):2803–2811, 2017.

30. Michelle E LaFrance, Melissa A Farrow, Ramyavardhanee Chandrasekaran, Jinsong Sheng, Donald H Rubin, and D Borden Lacy. Identification of an epithelial cell receptor responsible for clostridium difficile tcdb-induced cytotoxicity. Proceedings of the National Academy of Sciences, page 201500791, 2015.

31. Lionel Le Bourhis, Emmanuel Martin, Isabelle Péguillet, Amélie Guihot, Nathalie Froux, Maxime Coré, Eva Lévy, Mathilde Dusseaux, Vanina Meyssonnier, Virginie Premel, et al. Antimicrobial activity of mucosal-associated invariant t cells. Nature Immunology, 11(8): 701, 2010.

32. Young-Nan Cho, Seung-Jung Kee, Tae-Jong Kim, Hye Mi Jin, Moon-Ju Kim, Hyun-Ju Jung, Ki-Jeong Park, Sung-Ji Lee, Shin-Seok Lee, Yong-Soo Kwon, et al. Mucosal-associated invariant t cell deficiency in systemic lupus erythematosus. The Journal of Immunology, page 1302701, 2014.

33. Hirohito Kita. Eosinophils: multifaceted biological properties and roles in health and disease. Immunological Reviews, 242(1):161–177, 2011.

34. Margarita Dominguez-Villar and David A Hafler. Regulatory t cells in autoimmune disease. Nature Immunology, page 1, 2018.

35. Trevor Hastie, Robert Tibshirani, and Martin Wainwright. Statistical learning with sparsity: the lasso and generalizations. CRC press, 2015.

36. Weijie Su, Małgorzata Bogdan, Emmanuel Candes, et al. False discoveries occur early on the lasso path. The Annals of Statistics, 45(5):2133–2150, 2017.

37. Benjamin Hofner, Luigi Boccuto, and Markus Göker. Controlling false discoveries in high-dimensional situations: boosting with stability selection. BMC bioinformatics, 16(1):144, 2015.

